# Segment Anything for Microscopy

**DOI:** 10.1101/2023.08.21.554208

**Authors:** Anwai Archit, Sushmita Nair, Nabeel Khalid, Paul Hilt, Vikas Rajashekar, Marei Freitag, Sagnik Gupta, Andreas Dengel, Sheraz Ahmed, Constantin Pape

## Abstract

We present Segment Anything for Microscopy, a tool for interactive and automatic segmentation and tracking of objects in multi-dimensional microscopy data. Our method is based on Segment Anything, a vision foundation model for image segmentation. We extend it by training specialized models for microscopy data that significantly improve segmentation quality for a wide range of imaging conditions. We also implement annotation tools for interactive (volumetric) segmentation and tracking, that speed up data annotation significantly compared to established tools. Our work constitutes the first application of vision foundation models to microscopy, laying the groundwork for solving image analysis problems in these domains with a small set of powerful deep learning architectures.

## Main

Identifying objects in microscopy images, such as cells and nuclei in light microscopy (LM) or cells and organelles in electron microscopy (EM) is one of the key tasks in image analysis for biology. The large variety of imaging modalities and different dimensionality (2d, 3d, time) make these identification tasks challenging and so far require different approaches for different applications. The relevant state-of-the-art methods are deep learning based and have in the past years significantly improved cell and nucleus segmentation in LM^1–3^, cell, neuron and organelle segmentation in EM^4–7^ and cell tracking in LM^8,9^. Most of these methods provide pretrained models and yield high quality results for new data similar to the model training sets. However, due to limited generalization capabilities of the underlying deep learning approaches, quality degrades for data dissimilar to the original training sets, see also Ma *et al.*^10^, and they can only be improved by retraining on new annotated data. Generating these annotations is time consuming, as it relies on manual pixel-level annotations. CellPose 2^11^ implements in-the-loop annotation and retraining to speed up this process for 2D segmentation, but relies on manual pixel-level correction and is thus very time consuming if the initial segmentation result is of low quality.

Vision foundation models have recently been introduced for image analysis tasks in natural images, echoing developments in language processing. These models are mostly based on vision transformers^12^ and are trained on very large datasets. They can be used as a flexible backbone for different downstream tasks. The first successful foundation model was CLIP^13^, which combines images and language, and underlies many generative image models^14^. More recently foundation models targeting segmentation have been introduced^15,16^. Among them Segment Anything^15^ (SAM), which was trained on a large labeled dataset and achieves impressive interactive segmentation performance for a wide range of image domains. Such foundation models have so far not been applied in microscopy, but their potential in this domain has already been identified^17^.

Here, we introduce Segment Anything for Microscopy, which we call micro_sam in the following. It extends SAM to improve models for microscopy data and enable interactive data annotation for multidimensional data. Our main contributions are:

● We implement a training method to finetune SAM on new datasets.
● We use this method to finetune SAM models on microscopy data and find an overall significant improvement compared to the default models.
● We implement napari-based^18^ tools for interactive data annotation for (volumetric) segmentation and tracking that also support in-the-loop finetuning.

Figure 1 shows a high-level overview of micro_sam and examples for improved segmentation results. Prior work has already investigated SAM for biomedical applications, for example in medical imaging^19^, histopathology^20^ and neuroimaging^21^. However, these studies were limited to the default SAM models and did not implement customization for their respective domains, which is crucial according to our findings.

Compared to established segmentation and tracking tools, micro_sam is more versatile because its pretrained models work for both LM and EM. It also supports 2d and volumetric segmentation as well as tracking in the same set of tools and significantly speeds up data annotation due to the interactive segmentation capabilities of SAM. We demonstrate this in three user studies where we find competitive performance with CellPose^11^ for cell segmentation and significantly improved performance compared to ilastik carving^22^ for volumetric segmentation and compared to Trackmate^9^ for tracking. Our contribution also shows the promise of vision foundation models to unify image analysis solutions in bioimaging. Our tool is available at https://github.com/computational-cell-analytics/micro-sam and documented at https://computational-cell-analytics.github.io/micro-sam/micro_sam.html.

## Results

We compare the default SAM models with our finetuned models on a variety of microscopy segmentation tasks. First, we study the finetuning method and its variations on the LiveCELL^23^ dataset. Then we train and evaluate generalist models for LM and EM. Finally, we introduce interactive annotation tools for (volumetric) segmentation and tracking based on napari^18^. We compare them to established tools in three user studies.

### Finetuning Segment Anything significantly improves segmentation performance on the LiveCELL dataset

In Kirilov *et al.*^15^, SAM is introduced as a model for interactive segmentation: it predicts an object mask based on point or bounding box inputs, see also Fig. 1 a for an example. The point annotations can be positive (part of the object) or negative (not part of the object). The model was trained on a very large dataset of natural images with segmentation annotations. The authors also introduce a method for automatic instance segmentation. They evaluate interactive and automatic segmentation on a wide range of tasks, including a microscopy dataset^24^. See <u>Methods</u> for an overview of the SAM functionality. The original microscopy experiment and our evaluation of the default SAM models show a remarkable generalization to microscopy, despite only natural images in the original training set. However, we noticed several shortcomings of this model for microscopy. For example, it segments clusters of cells as a single object as seen in Figure 1 b. To improve it for application in our domain, we implement an iterative training scheme to enable finetuning on new datasets. This approach is inspired by the training method described in the SAM publication^15^, which has so far not been made open source. See <u>Methods</u> for details on the training scheme. We first evaluate default and finetuned SAM models as well as different training strategies on LiveCELL^23^, the largest publicly available dataset for cell segmentation. Figure 2 a shows the performance with increasing number of point annotations (box plot), with box annotations (blue line) and with the automated instance segmentation (green line) for the default (left) and finetuned (right) model. The point and box annotations are derived from the ground-truth segmentation provided by LiveCELL. We finetune the largest SAM model (ViT-H) on the training split of LiveCELL and use the test split for the evaluation experiments. We evaluate the segmentation results using the segmentation accuracy metric^25^. We also indicate the performance of CellPose^11^ trained on LiveCELL (yellow line). See <u>Methods</u> for more details on inference and evaluation. The results show a significant improvement due to finetuning across all settings. After finetuning segmentation from point or box annotations performs better than CellPose. The automated segmentation significantly improves but is not on par with CellPose. We investigate different finetuning strategies in Figure 2 b, where we finetune only parts of the SAM model and leave the rest of its weights frozen. Here, we use a smaller version of the model (ViT-B), but otherwise follow the same experimental set-up as before. The results show that finetuning the image encoder (IE) has the biggest impact, and finetuning the complete model shows the best overall performance.

**Figure 1:**
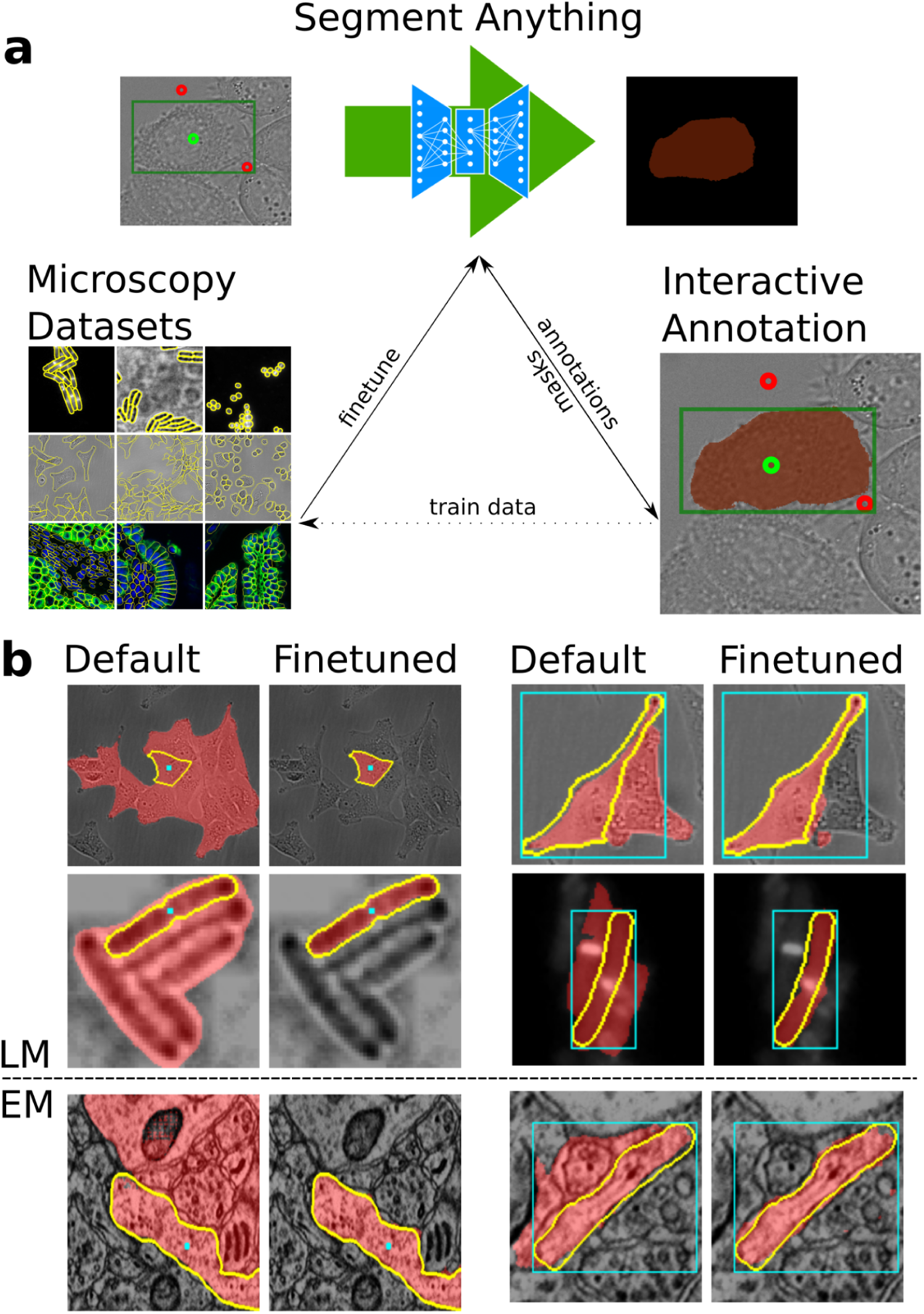
Overview of the Segment Anything for Microscopy contributions. **a.** We provide models that were customized for microscopy data by finetuning on large microscopy datasets. We implement napari-based interactive annotation tools for multi-dimensional microscopy data based on Segment Anything. These tools enable rapid data annotation for several applications and can be used to generate data for downstream analysis or for finetuning a custom model to further speed up annotation or to automatically process the data (not shown). **b.** Improvement of segmentation quality due to finetuning on light (top) and electron (bottom) microscopy data.

**Figure 2:**
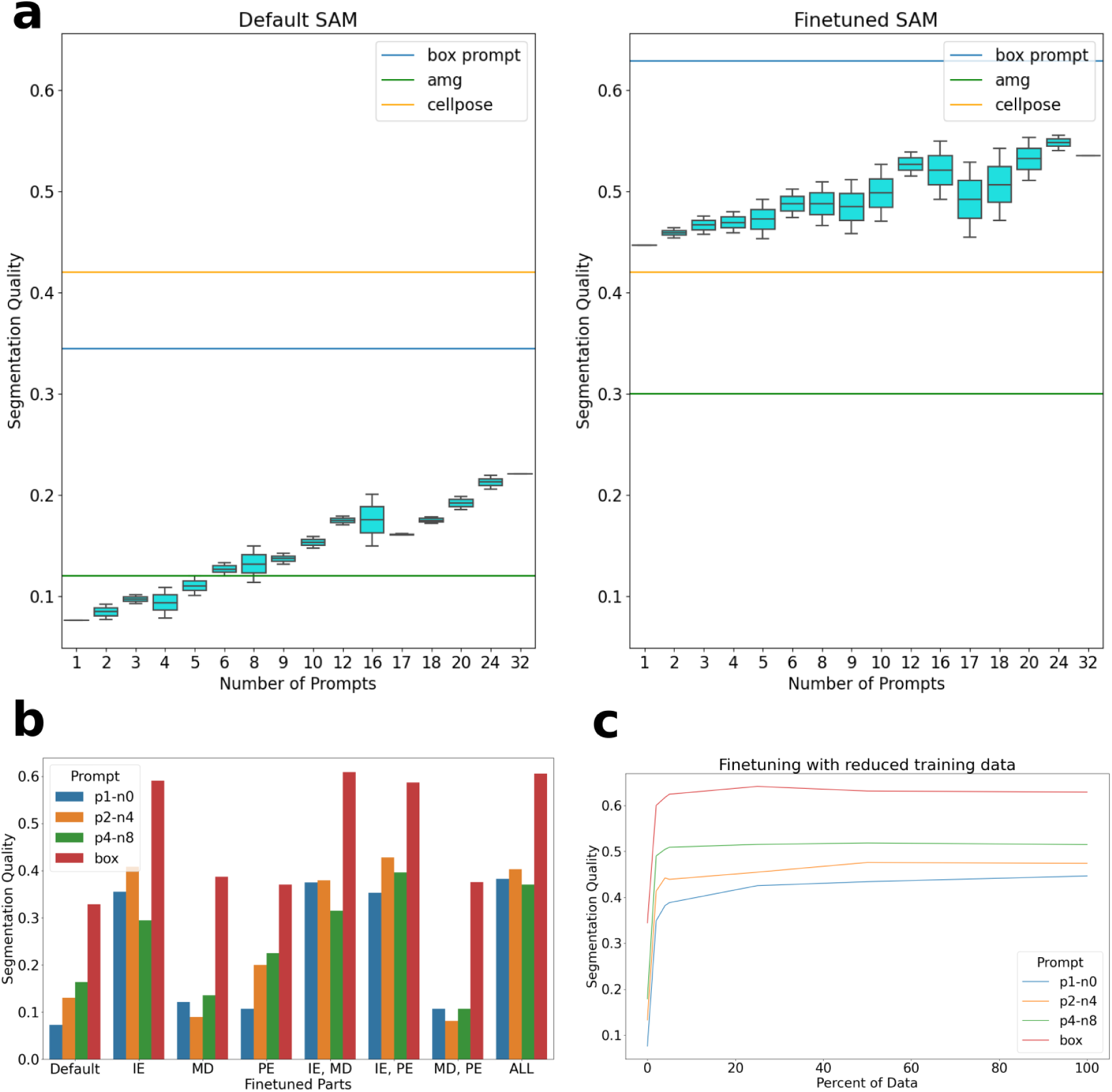
Results on LiveCELL^23^. **a** compares the default SAM model (left) with the finetuned version (right). The box plot shows the performance with increasing number of point annotations. We use different combinations of positive and negative points and report the total number on the x-axis. The lines indicate the performance of box annotations (blue), automatic instance segmentation (green) and CellPose (yellow). **b** compares different finetuning strategies where only parts of the model are finetuned. Here the label on the x-axis indicates which part(s) of the model are updated during training, IE stands for image encoder, MD for mask decoder and PE for prompt encoder. We evaluate box annotations (blue) and three different combinations of point annotations, where “p1-n0” means one positive and no negative point etc. **c** shows the segmentation quality for increasing size of the training dataset. The results of **a** and **c** are for the largest SAM model (ViT-H), **b** for the smallest one (ViT-B). Supp. Figure 1 shows an explanation of the model parts and results for the Vit-B version of SAM.

We also finetune the model with only a subset of the training data, using data splits defined in the LiveCELL publication. We use the same experimental set-up as before with the largest model. The results in Figure 1 c show that the majority of the quality improvement occurs already with the small training data fractions of 2, 4 and 5 %.

The results on LiveCELL offer the following important conclusions for further experiments:

1. Finetuning SAM significantly improves the segmentation quality for a given dataset.
2. Finetuning the full model yields the best results.
3. Significant improvements can be achieved for small training datasets.

The points 1 and 2 prescribe a recipe for training new “generalist” SAM models for the microscopy domain. Point 3 indicates that in-the-loop finetuning based on user annotations should be feasible. Unless stated otherwise, all further experiments will follow the same set-up for finetuning and evaluating models as in this section.

### Light microscopy model trained on diverse data generalizes to new experimental settings

Our next goal is to train a generalist model for LM. This model should improve segmentation performance for a wide range of segmentation tasks, so that it can be used as a replacement to the default SAM model for LM. While our previous experiments have shown that finetuning on data from a given imaging setting significantly improves specific performance, we have not yet shown improved generalization to other settings.

To train the generalist, we assemble a large and diverse training set based on published LM datasets: LiveCELL^23^, TissueNet^2^, DeepBacs^26^, Neurips Cell Segmentation^10^, Nucleus DSB^24^ and PlantSeg^27^. We train the generalist model on the combined training set, and for LiveCELL, TissueNet and DeepBacs also train specialist models. The top row in Figure 3 a shows the segmentation performance on these three datasets and compares default, specialist and generalist model. In all cases the evaluation is done on a test split that is not part of the training set for specialists and generalist. We see significant improvements of both specialists and generalist compared to the default model. The specialists overall show slightly better performance compared to the generalist, especially for DeepBacs, the smallest of the three datasets. However, this difference is negligible compared to the performance improvement compared to the default model. We also indicate the performance of a CellPose model as a baseline, using the *cyto* model, except for LiveCELL where the model trained on this dataset is used. Overall, the results show that training a generalist model on a superset of datasets does not lead to significant loss in performance for a given dataset.

**Figure 3:**
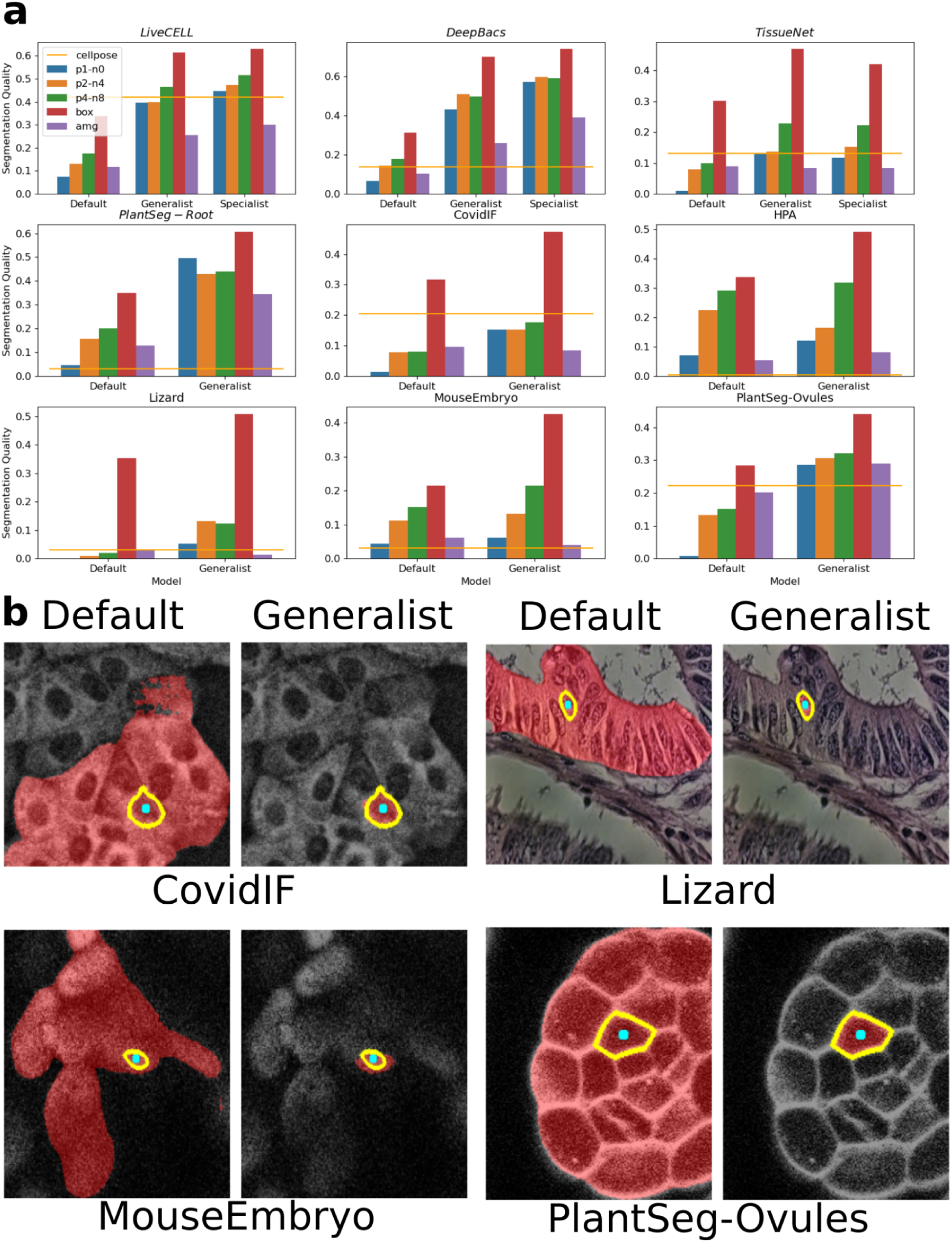
Generalist light microscopy model. **a** compares the performance of the default SAM model with our generalist and in some cases specialist models. Here, we use the same experimental settings as in Figure 2 b and c; in addition “amg” (purple) indicates the performance of automatic instance segmentation and the yellow line the performance of CellPose using the *cyto* model (except for LiveCELL where we use a more specific model).

To study whether the generalist model has improved generalization for different microscopy imaging settings we apply it to five new datasets, which come from settings not represented in the training set. We choose the datasets CovidIF^28^ containing immunofluorescence data, HPA^29^ containing fluorescence images of human cells, Lizard^30^ containing histopathology data, mouse-embryo^31^ containing nuclei imaged in light-sheet and plantseg-ovules^27^ containing cells imaged in confocal. The comparison of the default and generalist model as well as the CellPose *cyto* model are shown in the middle and bottom row of Figure 3 a. The improvement of the generalist over the default model is significant for all datasets. The comparison to CellPose shows mixed results. In some cases only the box annotations, which contain the most information about the object to be segmented, improve over CellPose, in other cases all SAM settings including the automated instance segmentation outperform it. Figure 3 b shows some qualitative comparisons of default and generalist model. Supp. Figures 2 and 3 show more qualitative comparisons, Supp. Figure 5 shows the results based on the ViT-B model; the experiments presented here are based on ViT-H. More details on the LM datasets can be found in Supp. Table 1.

Overall, these experiments demonstrate that generalist SAM models for a given domain can significantly improve performance and the generalist model we train improves interactive and automatic segmentation performance for cell and nucleus segmentation in LM.

Datasets in italic are part of the training set (evaluated on a separate test split). **b** shows interactive segmentation results with default and generalist model. The cyan dot corresponds to the point annotation, the yellow outline to the object and the red overlay to the model prediction.

### Electron microscopy model improves segmentation performance for cells and neurites

We further investigate if training a generalist model for EM is feasible. This is more challenging compared to LM because in EM membrane bound structures are labeled unspecifically rather than having a specific stain for a cellular component^1^. Since these structures have a hierarchical composition, e.g. organelles inside of a cell or cellular compartment, training a generalist model is more challenging. Nevertheless, we proceed similarly to the LM case and assemble a large training set consisting of the publicly available datasets CREMI^32^, MitoEM^33^, AxonDeepSeg^34^ and PlatyEM^4^. This set contains annotations for cells, cilia, mitochondria, myelinated axons, neurites and nuclei imaged with different EM techniques. Using separate test splits we compare the performance of the generalist model trained to the default model for four datasets. The results in the top row of Figure 4 a show significant improvements for cells, neurites and nuclei, but a decrease in performance for mitochondria.

**Figure 4:**
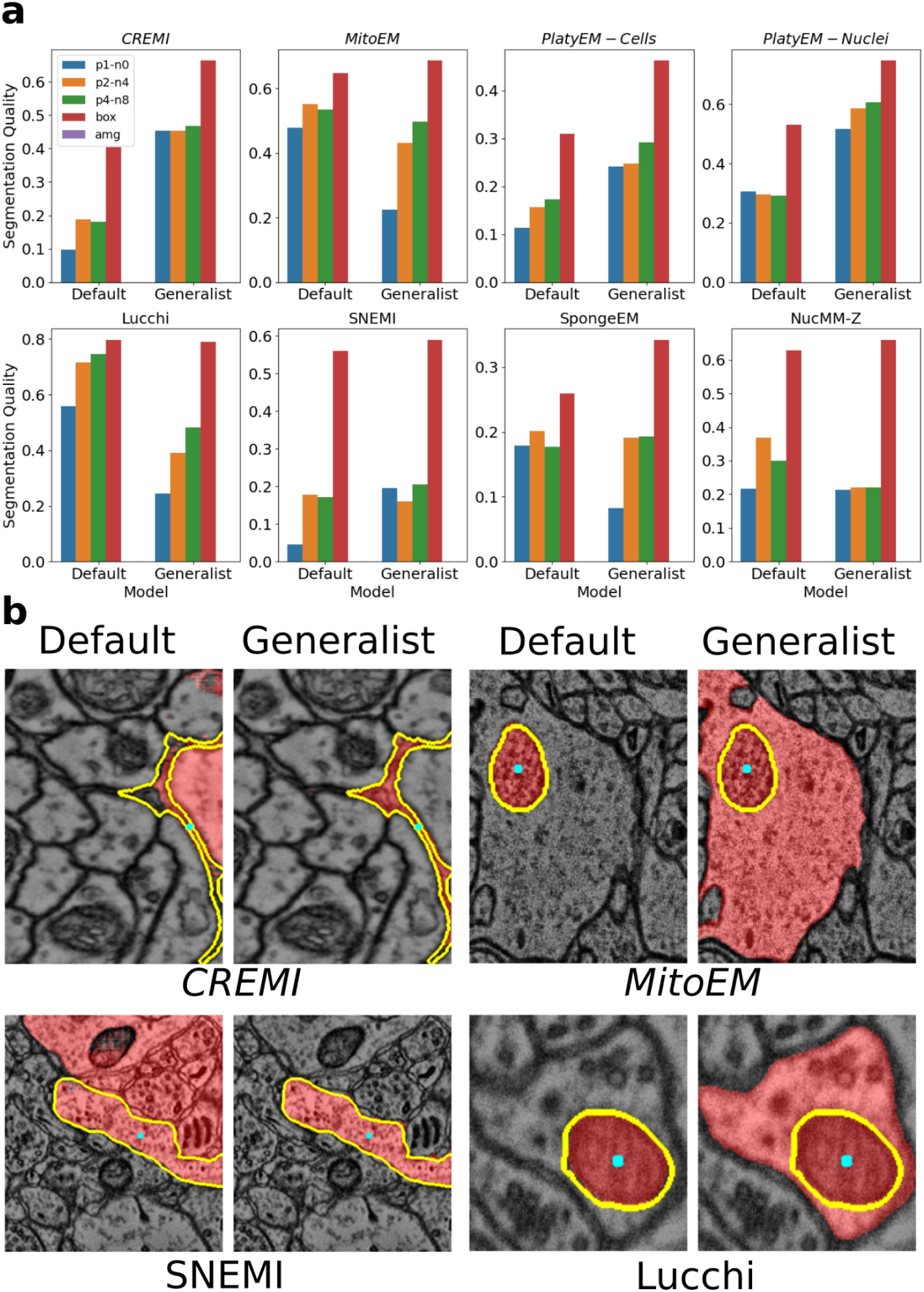
Electron microscopy models. **a** compares the performance of the default SAM model to our EM model. Datasets in italic are part of the training set (evaluation is done on separate test splits). We follow the same experimental set-up as before, but do not evaluate the automatic instance segmentation, since this is ill-defined given segmentation tasks for specific objects, and do not compare to a baseline method for similar reasons. **b** shows qualitative comparisons, using the same color coding as in Figure 3 b.

We also compare the generalist to the default model on four datasets with imaging conditions not present in the training set: Lucchi^35^ containing mitochondria imaged in FIBSEM, NucMM-Z^36^ containing nuclei imaged in low resolution SEM, SNEMI^37^ containing neurites imaged in SEM and SpongeEM^38^ containing cells and organelles imaged in FIBSEM. The results are shown in the bottom row of Figure 4. We see a similar trend as before: the performance improves for neurites, decreases for mitochondria and shows mixed results for other settings. Figure 4 b shows qualitative comparisons between the default and generalist model and more comparisons can be found in Supp. Figure 4. Supp. Figure 5 shows the result for ViT-B. Supp. Table 2 gives more details on all datasets.

These results show that training a generalist model for any structure in EM is not feasible given the current SAM architecture and training scheme. However, we see that an improved model can be trained for specific structures; in our case the EM model significantly improves performance for cells and neurites, which make up the largest part of the training set. Note that in the current state micro_sam can speed up annotation tasks in EM significantly as demonstrated in one of the user studies. However for structures other than cells, neurites or cellular compartments the default model will likely perform better than our finetuned model. We will provide dedicated EM models for mitochondria and other organelles soon.

### Interactive annotation tools based on Segment Anything speed up data annotation for diverse applications in microscopy

To make the interactive and automatic segmentation functionality offered by SAM available to microscopists and image analysts we implement three tools: one for 2d segmentation, one for volumetric segmentation and one for cell tracking. They are implemented using napari^18^, a python-based viewer for multi-dimensional image data. The 2d annotation tool supports interactive segmentation based on user-provided point and/or box annotations. It also offers automated instance segmentation functionality that can be used to generate an initial segmentation. Alternatively initial segmentations from other methods can be loaded by the tool. To enable smooth interactive annotation on a laptop without a GPU we extend the core SAM functionality to support precomputation and caching of image embeddings, tiled computation of embeddings and tiled interactive segmentation as well as efficient recomputation of the automatic instance segmentation given parameter changes.

For convenience we also implement an annotation tool for a series of input images. This tool can precompute the state for all images in a folder and then opens the segmentation annotator consecutively for these images, after the user saves the segmentation result for an image.

The volumetric annotation tool extends the interactive segmentation functionality to 3d data by projecting segmentation masks to adjacent slices and running SAM with annotations derived from the projected mask as inputs. It supports the same features as the 2d annotation tool, except for the automatic instance segmentation functionality.

The annotation tool for (cell) tracking works similarly by projecting masks to adjacent timeframes. It uses a linear motion model for projection, to account for the fact that cells often move in a given direction. This tool further supports annotation of division events, so that lineages of cells can be annotated. Otherwise it supports the same functionality as the 2d annotation tool, except for automatic segmentation.

The core functionality underlying these features can also be used programmatically from python, so that other developers can build upon our SAM extensions. See <u>Methods</u> for the implementation details of the extended functionality and the annotation tools. Supp. Figure 6 shows an overview of the user interface and Supp. Videos 1-3 show tutorials that explain the tool usage. The full documentation of our software can be found at https://computational-cell-analytics.github.io/micro-sam/micro_sam.html.

We study the annotation tools for three representative applications and compare them to established tools for the respective annotation tasks. For 2d annotation we study organoid segmentation in brightfield images. Growing organoids is a common experimental technique for studying tissue, e.g. in cancer research, and analysis based on organoid segmentation enables studying their growth and morphology, e.g. to compare the effect of different treatments. Here, we make use of an internal dataset for which 20 images have ground-truth segmentation masks and plenty more images without ground-truth exist. We compare different annotation approaches on this dataset and report the annotation time per object in Figure 5 a (left). We use manual annotation (“manual”), annotation and in-the-loop finetuning with CellPose 2 (“CellPose (default)”, “CellPose (finetuned)”) as well as different annotation approaches with micro_sam: interactive annotation without initial automated instance segmentation (“micro_sam (no amg)”), interactive annotation with automated instance segmentation (“micro_sam (default)”) and the same set-up with a model finetuned in-the-loop (“micro_sam (finetuned)”). We use the GUI provided by CellPose for annotation and finetuning of this method, starting from the *cyto* model. Here, we first annotate three images without updating the model and then finetune it twice on the previous annotations. For simplicity we only report the annotation times for the default model and the mean over annotation times after finetuning.

**Figure 5:**
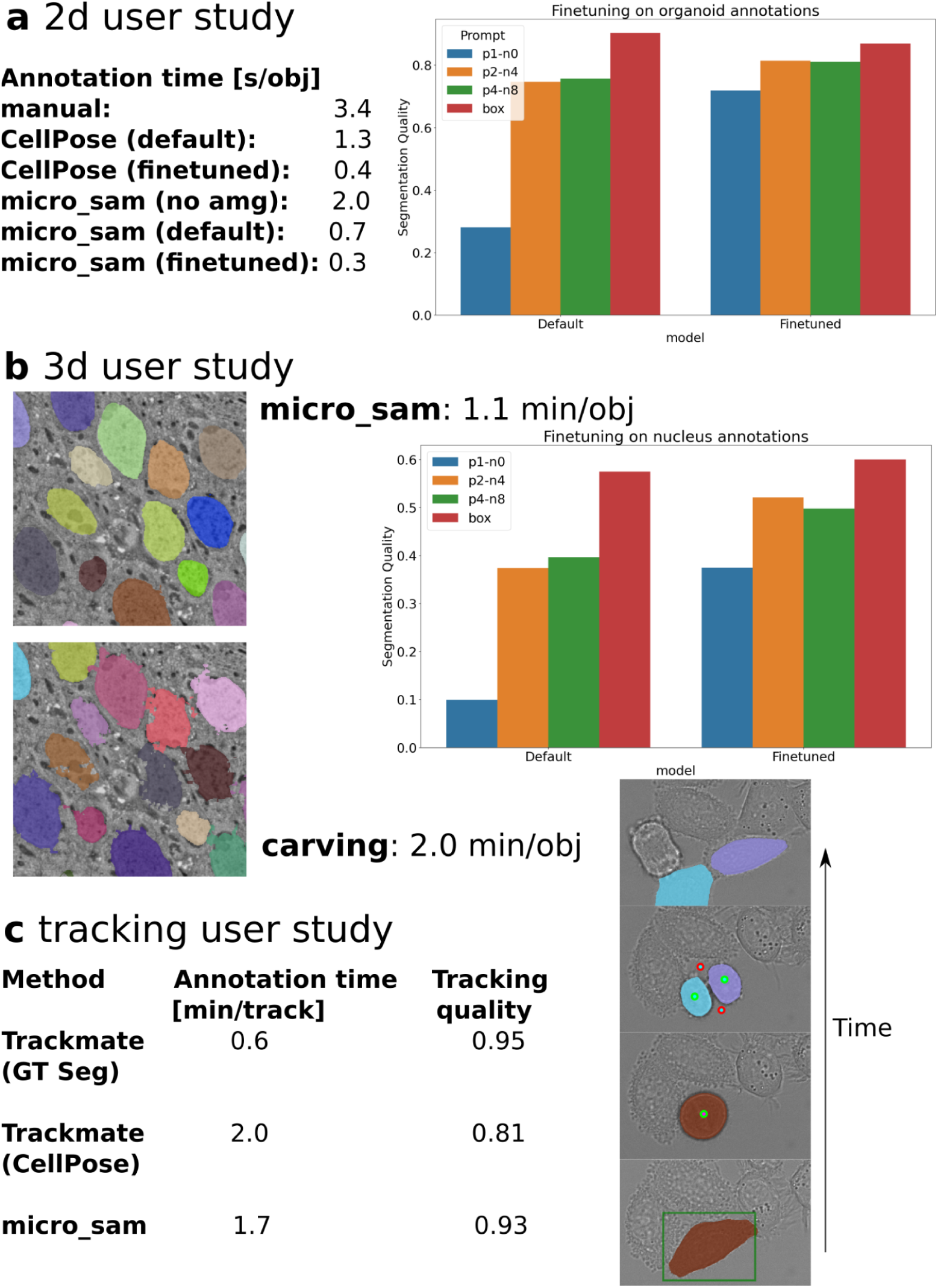
Annotation tool user studies. **a** Segmentation of organoids imaged in brightfield. Table on lists annotation times per object for variants of our tool, CellPose, and manual annotation. Plot on the right compares performance of the default SAM model with the model finetuned on user annotations. **b** Comparison of ilastik carving and 3d annotation with micro_sam. Images show a slice of the EM data with segmentation results from both tools overlayed (results are volumetric for both) and the annotation time per object is given. Plot on the right compares performance of the default SAM model with the model finetuned on user annotations. **c** comparison of Trackmate and micro_sam. Table lists annotation times and quality, the figure shows an example of a cell lineage reconstructed with micro_sam, including annotations.

For micro_sam we use the default SAM ViT-H model and first compare annotation times with and without starting from automated instance segmentation, where we find a clear advantage for the former. We then finetune the model on the segmentation results from interactive segmentation and also report the annotation times (using automatic instance segmentation) for this model. We see that all semi-automatic approaches perform significantly better than manual annotation. When using the automated instance segmentation micro_sam has an advantage over CellPose before finetuning due to its interactive segmentation capability. After finetuning, both methods perform comparatively. In this setting most organoids are correctly segmented automatically, and the advantage due to interactive segmentation becomes less relevant. We found that all approaches yield comparably high segmentation quality and do not evaluate it further. In addition we evaluate the quantitative performance of the default and finetuned SAM model in Figure 5 a (right), using data with labels that were not part of the data we have annotated on. We see a significant improvement in the performance for single point annotations and moderate improvements for the other settings. This improvement translates into the significant annotation speed up because the single point prompt setting is used by the automatic instance segmentation method and thus significantly improves its results.

For the 3d annotation tool we study nucleus segmentation in volume EM data, using an internal dataset of a fruitfly larva, for which we also have ground-truth annotations for several small blocks. Segmentation of nuclei or other large organelles in volume EM is an important task for analyzing cellular morphology and differentiating cell types based on phenotypic criteria^4^. Here, we compare interactively segmenting the nuclei with micro_sam and with ilastik carving^22^.

Carving uses a seeded graph watershed to segment objects in 3d from user annotations. This method is not based on deep learning, but is still one of the most commonly used approaches for 3d segmentation and is used by recent deep learning approaches for volume EM to generate training data^39^. In micro_sam we annotate the data with the default ViT-H model. Figure 5 b (left) shows two example slices from the volumetric segmentations and gives the annotation times.

Annotation with micro_sam is almost twice as fast and yields results of higher quality; the masks obtained from carving do not adhere well to the nuclei in some cases because of the superpixels that underlie the graph watershed and that cannot be corrected by carving. We also evaluate how finetuning on the user generated results improves the quantitative performance (Figure 1 b, right), observing significant performance improvements in most settings. Hence in-the-loop finetuning could speed up data annotation further as in the first use-case, but was not further studied here.

Finally, we study the tracking annotation tool on a dataset of HeLA cells imaged with label-free microscopy from the Cell Tracking Challenge^40^. We have extended this dataset to provide ground-truth segmentation for all slices in addition to the tracking ground-truth provided by the challenge. Cell tracking is a common analysis task for analyzing the dynamic behavior of cell populations. Here, we compare annotations with micro_sam with the most recent version of Trackmate^9^ that has integrated support for deep learning based segmentation tools, including CellPose. Figure 1 c shows the results for three different approaches: using Trackmate with the ground-truth segmentation as input, using it with the CellPose segmentation and using micro_sam. We report the annotation time per track, corresponding to the time it takes to track a cell between its start and end frame (it may end due to a cell division, because it goes out of frame, dies, or because the time series ends) and the tracking quality, measured by the metric from the Cell Tracking Challenge^40^. Note that we reconstruct lineages (i.e. tracks and division of mother into daughter cells) with all tools, but chose to report the time per track instead of lineage for simplicity. Annotation with Trackmate is fast and accurate when starting from ground-truth segmentation. However, without access to it and relying on the CellPose segmentation the annotation time increases significantly and the tracking quality decreases. In comparison data annotation with micro_sam is slightly faster and yields results of significantly better quality.

## Discussion

We have introduced methods for finetuning Segment Anything for microscopy and have built tools for interactive data annotation and automatic instance segmentation based on it. Our quantitative experiments and user studies show that our contributions significantly speed up data annotation and automated processing for a diverse set of applications. Our contribution also marks one of the first applications of vision foundation models in microscopy image analysis. We expect future work to build on our contributions and extend the application of foundation models in microscopy to further improve object identification tasks and address more image analysis tasks.

We compare our method to established tools for segmentation and tracking and show competitive or significantly improved performance. However, we expect that further improvements towards usability and performance can be made by integrating parts of our methods with them instead. For example, our finetuned models and interactive segmentation could be integrated within the CellPose GUI. The interactive segmentation and tracking functionality could also be integrated in Trackmate, where it could improve over the available segmentation options and be used to interactively correct tracks that were wrongly identified by automated tracking. Multiple ilastik workflows could also benefit from an integration, for example by implementing an advanced object detection workflow that makes use of the segmentation results from ilastik pixel classification. To support integration with other tools we have implemented the extensions to SAM’s functionality in a modular fashion, so that they can be used independently from our napari-based annotation tools.

We plan to improve and extend our methods and tools across several dimensions. In the near future we plan to train further models for bioimaging applications, for example a model specialized for mitochondria and other organelles in EM. We are also looking into recent developments that claim similar performance to the original SAM models with smaller and faster architectures^41^ that could speed up preprocessing times and interactive annotation in our tools. Furthermore, our current finetuning methodology relies on updating the full model weights. This has the disadvantage that it requires a GPU and can be quite time consuming, limiting in-the-loop finetuning to users with a powerful enough machine or access to on-premise or cloud GPUs. We envision that smaller models and advanced finetuning approaches like LoRA^42^ will make finetuning on consumer hardware feasible.

Finally, we think that the automatic instance segmentation functionality of SAM can be further improved by incorporating ideas from embedding based segmentation^43^. Such developments could also enable semantically aware instance segmentation, which would benefit domains like EM where structures of many different types are visible in the image.

## Methods

### Segment Anything

SAM is a vision foundation model for interactive segmentation. It was introduced in Kirilov *et al.*^15^. Here, we briefly summarize the main functionality of this method. It solves interactive segmentation tasks by predicting object masks based on an input image and annotations for a given object. The annotations can be either bounding boxes, points (positive and/or negative) or low-resolution masks. The publication also describes segmentation based on text annotations, but the available version of the model does not yet include this feature. For a new image, the model predicts an image embedding, which corresponds to a vector per pixel in a downscaled representation of the input, with the image encoder. The image encoder is a vision transformer^12^ and SAM comes in three variants with different sizes of this encoder, using the ViT-B, ViT-L or ViT-H architectures (ordered by increasing model size). The image encoder contains the majority of parameters of SAM and only gets the image data as input. It only has to be computed once per image, enabling fast recomputation of the segmentation mask if the annotations change and thus interactive segmentation. The other parts of the model are the prompt encoders that encode the box, point and mask annotation inputs and the mask decoder that predicts the object mask and intersection over union (IOU) score based on the image embedding and the encoded annotations. The IOU score corresponds to an estimate for the mask quality. To deal with the ambiguity of a single point annotation, which could refer both to an object or a part thereof, SAM predicts three different masks for this case. See also Supp. Figure 1 a for an overview of the SAM architecture.

The SAM model is trained on a large labeled dataset of natural images that is constructed iteratively by annotators that correct the outputs of a SAM model trained on a previous version of this dataset. The model is then evaluated on a broad range of segmentation tasks and shows remarkable generalization performance to image data from different domains. SAM requires per object annotations to predict segmentation masks. The authors also implement a method for automatic instance segmentation. It covers the input image with a grid of points and predicts masks for all points. The predicted masks are post-processed through several steps to retain only high quality predictions. This involves filtering out masks with a low IOU prediction, and masks with a low stability score, which is computed based on the change of the masks when thresholded at different logit values. In addition non-maximum-suppression is applied to remove overlapping predictions.

### Training

To finetune SAM models on custom data we implement an iterative training scheme similar to the training procedure described by Kirilov *et al.*^15^. Note that this part of their code has so far not been released. The training scheme requires image data and corresponding ground-truth segmentations for the objects of interest as input.

As is common when training deep neural networks, we iterate over the complete training dataset several times in so called epochs. In a single iteration we sample a minibatch, corresponding to multiple images and corresponding ground-truth, present the images to the network, compute the loss between predictions and ground-truth and update the network weights via backpropagation and gradient descent. Compared to usual training approaches a single iteration in our scheme is more complex and follows multiple steps:

1. Sample a minibatch containing input images and ground-truth from the training set.
2. Sample a fixed number of objects from the ground-truth. Training with all present objects would be too memory intensive.
3. We then perform the following steps for all sampled objects in a batched fashion:
  a. Sample either a random point from the object, which is used as a positive input point, or use the bounding box of the object as input.
  b. Predict the mask and expected IOU value for the given input with SAM. If the model is presented with only a single point annotation, predict three output masks, otherwise only predict a single output mask. See the discussion about ambiguity above for the motivation of this approach.
  c. Compute the loss between the predicted and ground-truth object as well as the loss between the estimated and true IOU between predicted object and ground-truth.
  d. Sample two new points, a positive one where the model predicted background but where there should be foreground (according to the ground-truth), and a negative one where settings are reversed. In case such points cannot be sampled because there is no region with missing foreground predictions or with versa we just sample a random positive / negative point.
  e. Present the combined annotations from the previous steps, i.e. all points sampled so far and the box annotation if used in the first step, as well as the mask prediction from the previous step as annotation inputs to the model.
  f. Compute the mask and IOU loss for the current predictions.
  g. Steps d, e and f are repeated for a fixed number of times, all losses are accumulated; backpropagation and gradient descent are performed based on the average loss over all steps
4. For efficiency reasons the image embeddings are computed once per image before step 3. Note that the image embeddings only depend on the image data, not on the input annotations.

The goal of this training scheme is for the model to iteratively improve segmentation masks and provide a valid mask output for any kind of input annotations. We have also experimented with simpler training schemes that did not make use of multiple annotation steps and instead relied on sampling box and and a fixed number of point annotations from the ground-truth. However, we found that this approach leads to significantly worse results.

For validation we rely on this simpler approach to generate input annotations and use the Dice score of prediction and ground-truth masks as metric. All experiments reported in this manuscript rely on finetuning the default SAM weights provided by the SAM publication. Our training method could also be used to train a model from randomly initialized weights. However, we expect this approach to significantly increase training times and thus did not pursue it.

We use the following settings and hyperparameter for training:

● We use a batch size of two. I.e. two input images and corresponding ground-truth are sampled per batch.
● We use the Dice loss to compare ground-truth objects and mask predictions.
● We use the L2 loss to compare true and predicted IOU scores.
● We use the ADAM optimizer^44^ with an initial learning rate of 10^-5^.
● We lower the learning rate when the validation metric plateaus (ReduceLROnPlateau).
● We train the models for 100 thousand iterations, except for:
  ○ The models where only parts of the architecture are updated (Figure 1 b). Since we trained SAM with the smaller ViT-b architecture in these cases we only trained for 25 thousand iterations.
  ○ The finetuned models in the user-study, which were only trained for 10 thousand iterations due to the limited training data and in-the-loop setting.
● All models were trained on a A100 GPU with 80 GB of VRAM. Training a model for 100 thousand iterations took about two days.

For the implementation we reuse the code from Kirilov *et al.* wherever possible and implement the additional training logic with PyTorch and torch-em^2^, a PyTorch based library for deep learning applied to microscopy also developed by us.

### Inference and evaluation

To quantitatively evaluate the interactive segmentation performance of SAM models we derive input annotations from ground-truth. Here, we don’t use an iterative approach, but either use object bounding boxes or sample a fixed number of positive and negative points per object.

When sampling points we always use the eccentricity center of an object as the first point, to mimic manual annotation, which usually places the first point near the center of an object. All other points are randomly sampled from the object mask (positive points) or from outside of it (negative points). For reproducibility we compute and store all point annotations for a given dataset and then use the same annotations for all models that are evaluated on it.

When evaluating the automated instance segmentation we found that it was crucial to also optimize two of the hyperparameters: the IOU and stability thresholds that are used for filtering out low quality predictions, see also the first <u>Methods</u> section. While the default settings work well for the original SAM models they have to be lowered for the finetuned models. Presumably this is due to the fact that these models are better calibrated to the actual prediction quality for objects in microscopy, which is lower compared to natural images. To efficiently perform a grid search we precompute the predicted object masks and then evaluate the hyperparameter ranges to be tested. See the next section for details. The parameter search is performed on a separate validation set, and the best setting found is applied to the test set.

To evaluate instance segmentation results we make use of the mean segmentation accuracy averaged over object matching thresholds in the range from 0.5 to 0.95 with increments of 0.05. This metric was introduced in Everingham *et al.*^25^ and has been popularized for microscopy by the DSB Nucleus Segmentation Challenge^24^. Confusingly it is sometimes referred to as mean average precision in the context of microscopy, although this term refers to a different metric. See Hirling *et al.*^45^ for an in-depth discussion of the interpretation of segmentation metrics and their history in microscopy segmentation. To evaluate the quality of tracking results we make use of the tracking metric introduced by the Cell Tracking Challenge^40^.

### Interactive annotation tools & python library

In addition to training and inference logic, we extend the functionality of SAM in four major ways to enable efficient interactive segmentation:

● We implement precomputation of the image embeddings. This step takes the bulk of computation time and is independent of the user-provided annotations. It can thus be precomputed for an image so that subsequent interactive annotation becomes possible. Typical processing times on a laptop without GPU are about a minute for the embedding computation (per image) and less than a second for interactive segmentation. We also support caching the precomputed embeddings to file, so that they do not have to be recomputed upon restarting the annotation tool for a given input. This is of particular importance for volumetric or time series data. Image embeddings can also be precomputed on a separate resource with a GPU (*e.g.* local workstation, on-premise computer cluster or cloud) and then be copied to the laptop used for annotation.
● We implement tiled computation for embeddings and interactive segmentation. When using this feature the embeddings are computed for overlapping tiles of the input image. In interactive segmentation the tile that best matches the current annotations is chosen and the segmentation is computed for it; the overlap ensures that objects that are part of multiple tiles can be segmented and the overlap size has to be chosen accordingly. Tiled embeddings are also precomputed and can be saved to file. Tiling enables interactive annotation for large images that could not be processed with SAM “en block”. We also extend automatic instance segmentation to support tiled embeddings.
● For automatic instance segmentation we separate the computationally expensive process of predicting masks from the post-processing of the predicted masks. This separation enables precomputation of the state for automatic instance segmentation, which can be stored together with the embeddings. This way users can also interactively change the most important post-processing hyperparameters, in particular the confidence threshold for retaining predicted mask and minimal size of mask, to find the best automatic segmentation settings for their images. This feature is also used to efficiently perform the grid search mentioned in the previous section.
● We extend SAM’s segmentation functionality to multiple dimensions (volumetric or 2d + time) by projecting input annotations derived from a given object to adjacent slices / frames. We can derive (low-resolution) input masks, the bounding box and point annotations and can present SAM with any combination of these input annotations. In practice we found that the combination of box and mask annotation usually work best and choose it as the default. The projection can optionally be done based on a linear motion model, which can be beneficial for tracking objects that follow a directed movement. Objects are segmented / tracked through the volume / timeseries by consecutive projection to adjacent slices / frames. The IOU between slices / frames can be used as a stopping criterion to prevent following objects that are wrongly segmented or not present anymore (e.g. because the structure being segmented has stopped or the cell being tracked leaves the frame). This functionality also supports interactive correction: after adding annotations in a slice / frame the whole object / track will be recomputed accordingly.

The data annotation tools are implemented using napari and use existing napari functionality whenever possible. We use the napari *label layer* to represent object masks from interactive and automatic segmentation predictions. Hence all features of this layer type, such as importing segmentations from file, manually editing objects in the layer or saving the layer to an image file, are supported. We use the *point layer* to represent point annotations and the *shape layer* for box annotations. All additional GUI elements of our tools are generated with magicgui^46^, which generates the elements from type-annotated function signatures. All annotation tools support interactive instance segmentation via point and box prompts. The interactive segmentation functionality is implemented with four layers: the *prompt* and *box prompt* layers are used to create the point and box annotations for interactive segmentation. The *current_object* layer contains the objects that are currently being annotated, and the *commited_objects* layer contains objects that have already been annotated. Once the user is done with the current object(s) they press a button to move it (them) to *commited_objects*. In the case of the tracking tool these two layers are called *current_track* and *committed_tracks*. All tools precompute the image embeddings for the complete image data, so that only prompt encoder and mask decoder have to be reapplied when the annotations change, enabling response times of below one second on a consumer laptop. They also all support tiled embeddings to enable annotation of large image data.

The 2d annotation tool further supports automatic instance segmentation. Its predictions are stored in a separate layer and the state for automatic instance segmentation is precomputed to enable interactively changing hyperparameters as described above. When using box annotations the 2d annotation tool can segment multiple objects at a time, one per box. When using point annotations or a mix of box and point annotations only a single object can be segmented at a time. We also implement an annotation tool for a series of images, e.g. for multiple images stored in a folder. This tool precomputes the state for all images, and then enables the users to iteratively annotate them, automatically saving the segmentation results per image to a predefined folder.

The 3d annotation tool uses the interactive multi-dimensional segmentation functionality described above to implement volumetric segmentation. It does not use a motion model for projection. It has similar features as the 2d annotator, but does not support automatic instance segmentation and can only segment one object at a time regardless of the type of annotation used.

The tracking annotation tool also uses the multidimensional segmentation functionality, with optional motion model. It adds support for dividing objects by extending the point annotations by properties to mark division events and to differentiate separate objects (mother and daughter cells) within a lineage. Otherwise it supports similar features to the 2d annotator, without support for automatic segmentation and restricted to tracking a single lineage at a time.

All our software is implemented within a single python library, also including the training and inference functionality described in the previous two sections. To enable programmatic use we implement this library in a modular fashion, in particular to enable using most of the functionality without requiring importing napari and starting a GUI.

## Code and Data Availability

Our software is available on github under a permissive open source license at https://github.com/computational-cell-analytics/micro-sam. At the submission of this manuscript the software version is 0.2.2. We make use of publicly available datasets for most experiments. They are listed in Supp. Table 1 and 2. We use internal datasets for the 2d and 3d annotation user studies. These datasets will be made available upon request.

## Acknowledgements

We would like to express their gratitude to Sartorius AG for support of this research through the Quantitative Cell Analytics Initiative (QuCellAI) and would also like to extend our thanks to all partners involved in the initiative for their contributions and valuable insights. We also gratefully acknowledge the computing time granted by the Resource Allocation Board and provided on the supercomputer Lise and Emmy at NHR@ZIB and NHR@Göttingen as part of the NHR infrastructure. The calculations for this research were conducted with computing resources under the project nim00007. The work of Sushmita Nair for this manuscript was funded by the Deutsche Forschungsgemeinschaft (DFG, German Research Foundation) under Germany’s Excellence Strategy – EXC 2067/1-390729940.

We would also like to thank Genevieve Buckly for contributions to our software, Timo Lüddecke for helpful discussions on designing training objectives for SAM as well as Sebastian von Haaren for help with creating the tutorial videos. In addition we would like to acknowledge the groups of Günter Schneider and Marian Grade at University Medicine Göttignen as well as Michael Pankratz at University Bonn for providing data for the user studies.

## Author Contributions

CP, AA, SA and NK conceptualized the work. AA, CP and PH implemented the software. AA, CP, SN, VR, NK, SG and MF performed experiments. AA and CP visualized the results. CP and AA drafted the manuscript. All co-authors reviewed and edited the final manuscript.

## Supplementary Information

### Supplementary Videos

<u>Supplementary Video 1:</u> Tutorial for the 2d annotation tool. The video is available at https://youtu.be/ket7bDUP9tI.

<u>Supplementary Video 2:</u> Tutorial for the 3d annotation tool. The video is available at https://youtu.be/PEy9-rTCdS4.

<u>Supplementary Video 3:</u> Tutorial for the tracking annotation tool. The video is available at https://youtu.be/Xi5pRWMO6_w.

### Supplementary Tables

**Supplementary Table 1:** Overview of LM datasets.

| Dataset | Image Modality | Annotations | Source | Used for |
| --- | --- | --- | --- | --- |
| LiveCELL | Phase Contrast | Cells | Edlund *et al.*<sup>23</sup> | Generalist Training and Evaluation |
| DeepBacs | Various | Bacteria | Spahn *et al.*<sup>26</sup> | Generalist Training and Evaluation |
| TissueNet | Whole Tissue | Cells | Greenwald *et al.*<sup>2</sup> | Generalist Training and Evaluation |
| PlantSeg-Root | Lightsheet | Cells | Wolny *et al.*<sup>27</sup> | Generalist Training and Evaluation |
| DSB Nuclei | Fluorescence | Nuclei | Caicedo *et al.*<sup>24</sup> | Generalist Training |
| Neurips CellSeg | Various | Cells | Ma *et al.*<sup>10</sup> | Generalist Training |
| CovidIF | Immuno-fluorescence | Cells | Pape *et al.*<sup>28</sup> | Generalist Evaluation |
| HPA | Fluorescence | Cells | Ouyang *et al.*<sup>29</sup> | Generalist Evaluation |
| Lizard | Histopathology | Nuclei | Graham *et al.*<sup>30</sup> | Generalist Evaluation |
| MouseEmbryo | Lightsheet | Nuclei | Bondarenko *et al.*<sup>31</sup> | Generalist Evaluation |
| PlantSeg-Ovules | Confocal | Cells | Wolny *et al.*<sup>27</sup> | Generalist Evaluation |
| CTC HeLA | Label-free | Cells | Ulman *et al.*<sup>40</sup> | User study |

**Supplementary Table 2:**
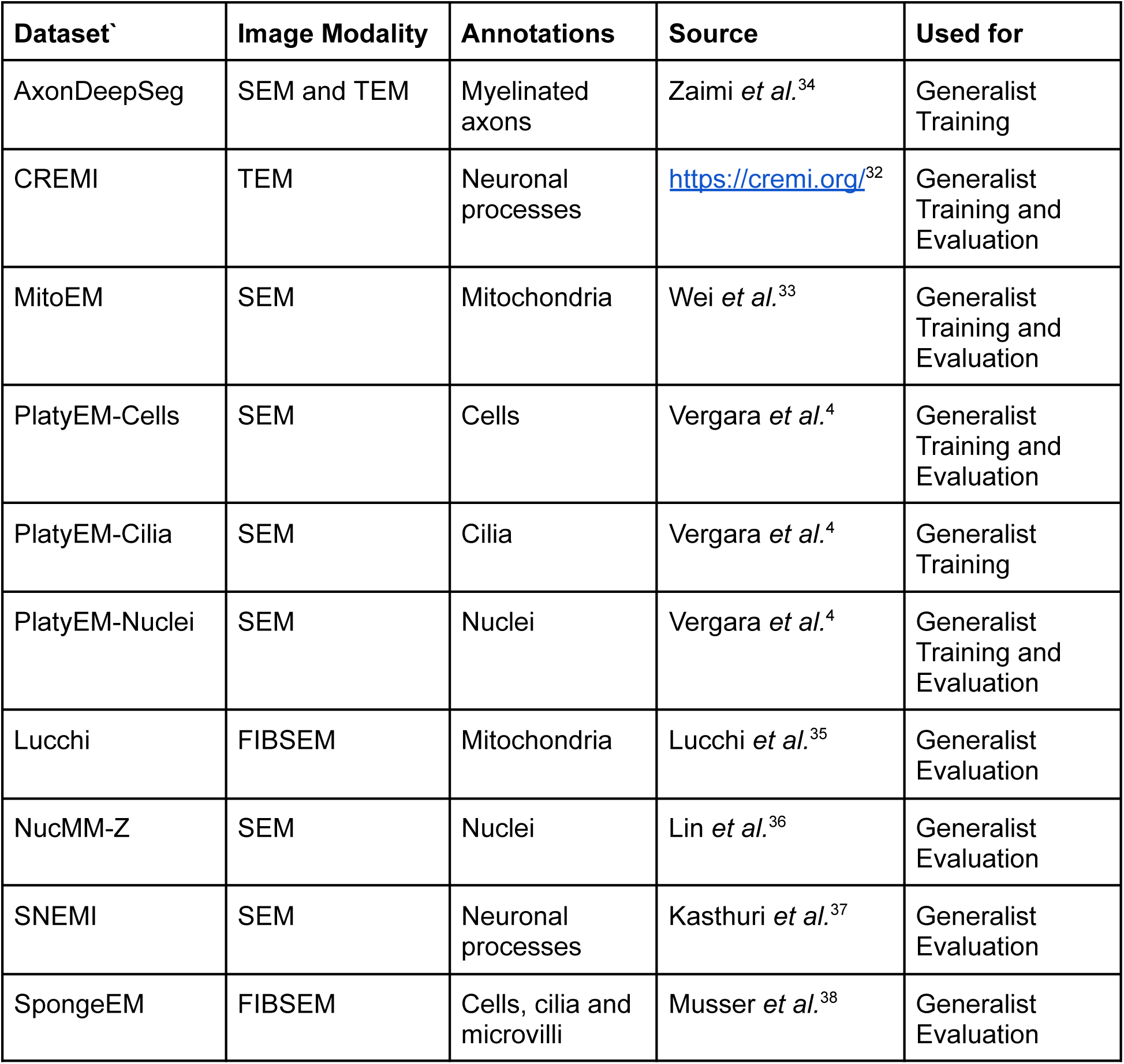
Overview of EM datasets.

| **Dataset** | **Image Modality** | **Annotations** | **Source** | **Used for** |
| --- | --- | --- | --- | --- |
| AxonDeepSeg | SEM and TEM | Myelinated axons | Zaimi *et al.*<sup>34</sup> | Generalist Training |
| CREMI | TEM | Neuronal processes | https://cremi.org/<sup>32</sup> | Generalist Training and Evaluation |
| MitoEM | SEM | Mitochondria | Wei *et al.*<sup>33</sup> | Generalist Training and Evaluation |
| PlatyEM-Cells | SEM | Cells | Vergara *et al.*<sup>4</sup> | Generalist Training and Evaluation |
| PlatyEM-Cilia | SEM | Cilia | Vergara *et al.*<sup>4</sup> | Generalist Training |
| PlatyEM-Nuclei | SEM | Nuclei | Vergara *et al.*<sup>4</sup> | Generalist Training and Evaluation |
| Lucchi | FIBSEM | Mitochondria | Lucchi *et al.*<sup>35</sup> | Generalist Evaluation |
| NucMM-Z | SEM | Nuclei | Lin *et al.*<sup>36</sup> | Generalist Evaluation |
| SNEMI | SEM | Neuronal processes | Kasthuri *et al.*<sup>37</sup> | Generalist Evaluation |
| SpongeEM | FIBSEM | Cells, cilia and microvilli | Musser *et al.*<sup>38</sup> | Generalist Evaluation |

### Supplementary Figures

**Supplementary Figure 1:**
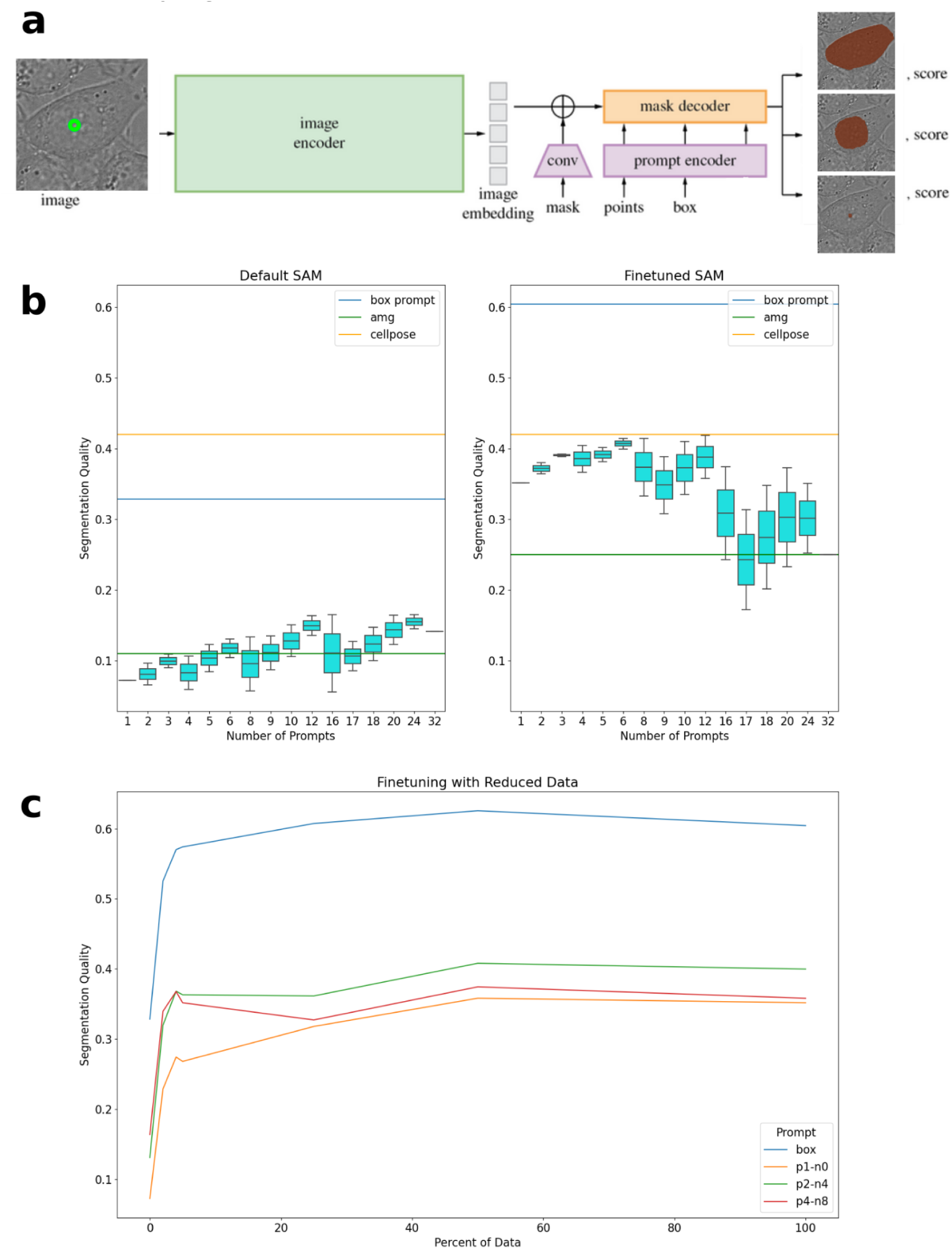
SAM architecture and ViT-B LiveCELL results. **a** the SAM model takes the image data and an annotation for the object to segment as input and outputs mask prediction(s) for the object. The architecture consists of the image encoder (IE) that computes an embedding for the image independent of the annotations, the prompt encoders (PE) that encode the mask, point and/or box annotations and the mask decoder (MD) that predicts the output mask and score. In the case of annotations with a single point the model predicts three potential output masks to deal with ambiguity. See for example the cell in the input image. The point annotation could refer to the cell, the nucleus or sub-texture of the nucleus. The score predicted by the model corresponds to the confidence of the model that the predicted mask is correct. This figure is adapted from Kiriliov *et al.*^15^. **b** SAM model with ViT-B evaluated for different annotation strategies on LiveCELL. Other than the ViT-B instead of ViT-H backend uses the same set-up as Figure 2 a. Unlike for the ViT-B model we see a decrease in performance with increasing number of point prompts for the finetuned model. **c** SAM model with ViT-B backend finetuned on subsets of the LiveCELL training data. We see the same overall trends as for ViT-H in Figure 2 c.

**Supplementary Figure 2:**
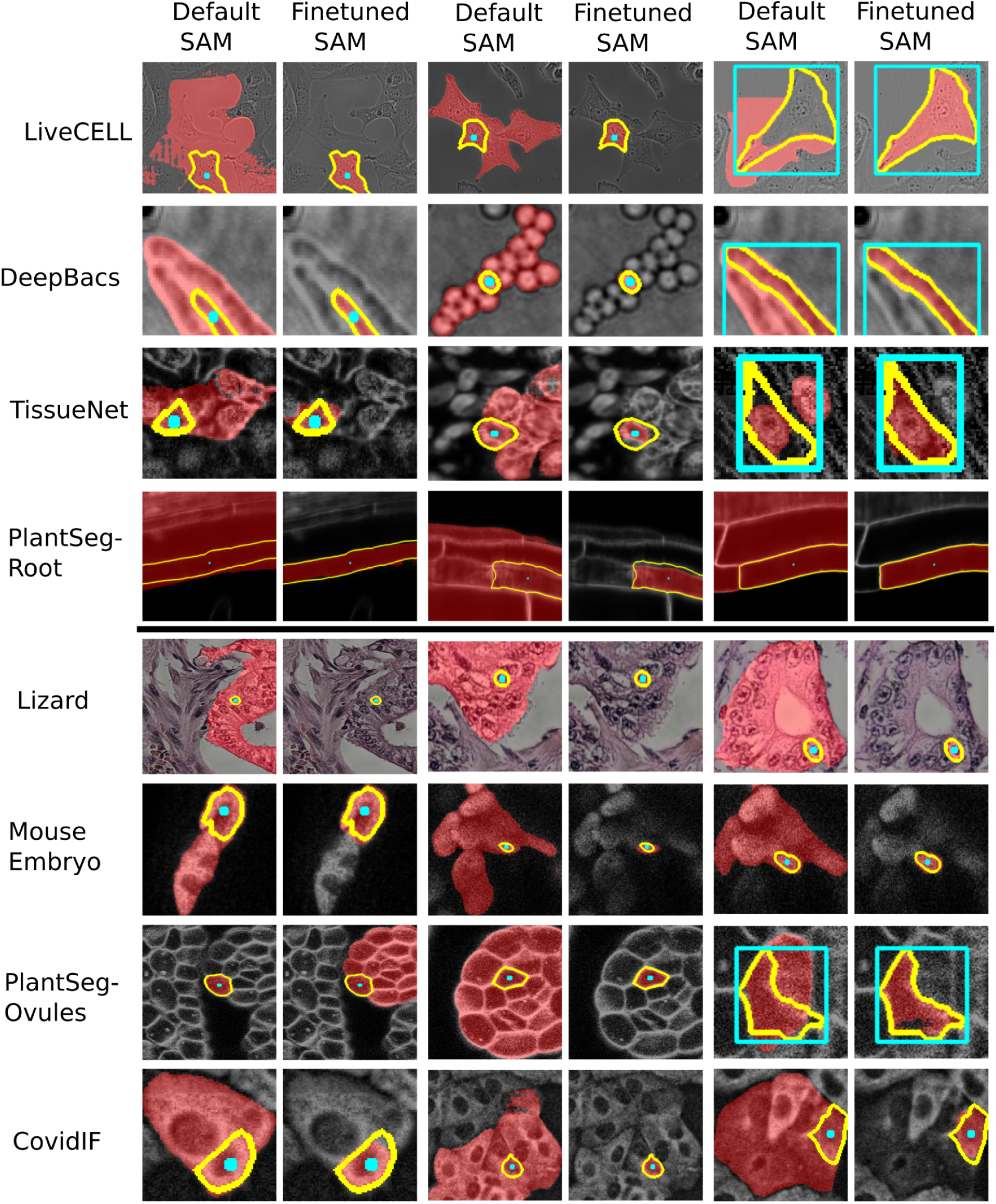
Qualitative comparison of LM generalist and default model. These images were generated by selecting objects with the highest improvement of IOU score of the finetuned compared to the default model for the given datasets. This approach shows that the biggest source of improvement for point annotations is the fact that the finetuned model only segments single objects in crowded situations, whereas the default model segments the whole local cluster of cells. For box annotations we see a better segmentation quality for objects of irregular shape and orientation.

**Supplementary Figure 3:**
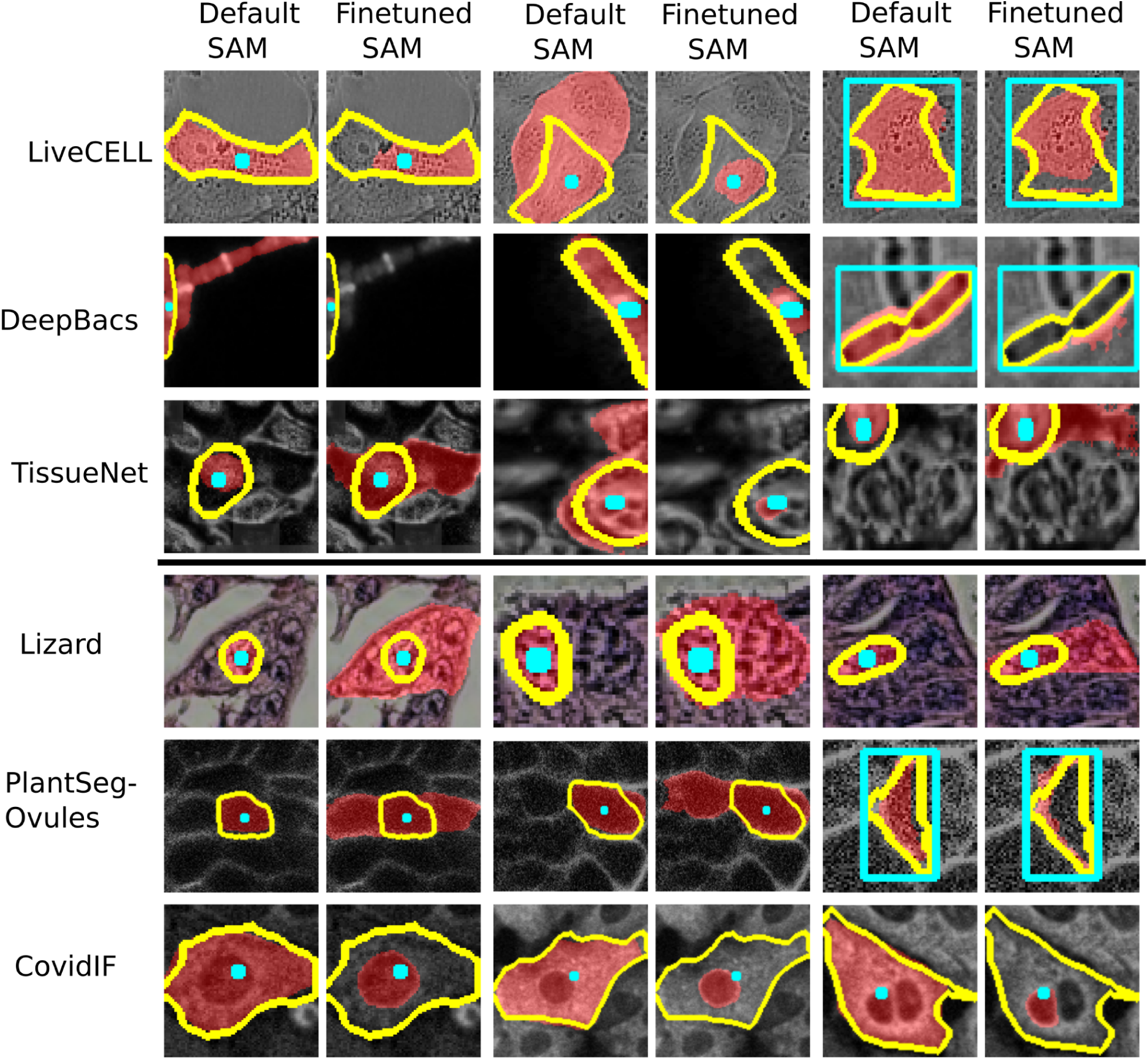
Qualitative comparison of LM generalist and default model. THis figure uses the opposite approach to Supp. Figure 2 and show the objects where the decrease in IOU is largest comparing the finetuned to default model. Here, we see a few different effects: in some cases the generalist model segments several nearby cells (proving an exception to the general behavior observed previously) for point annotations, in other cases the segmentation quality is lower given point or box annotations, sometimes segmenting smaller sub-structures. A clear systematic effect can be observed for CovidIF, where the generalist often segments only the nucleus, which is clearly discernible from the rest of the cell, rather than the full cell. Note that the overall segmentation quality for all these datasets is significantly higher for the generalist model as shown in Figure 3.

**Supplementary Figure 4:**
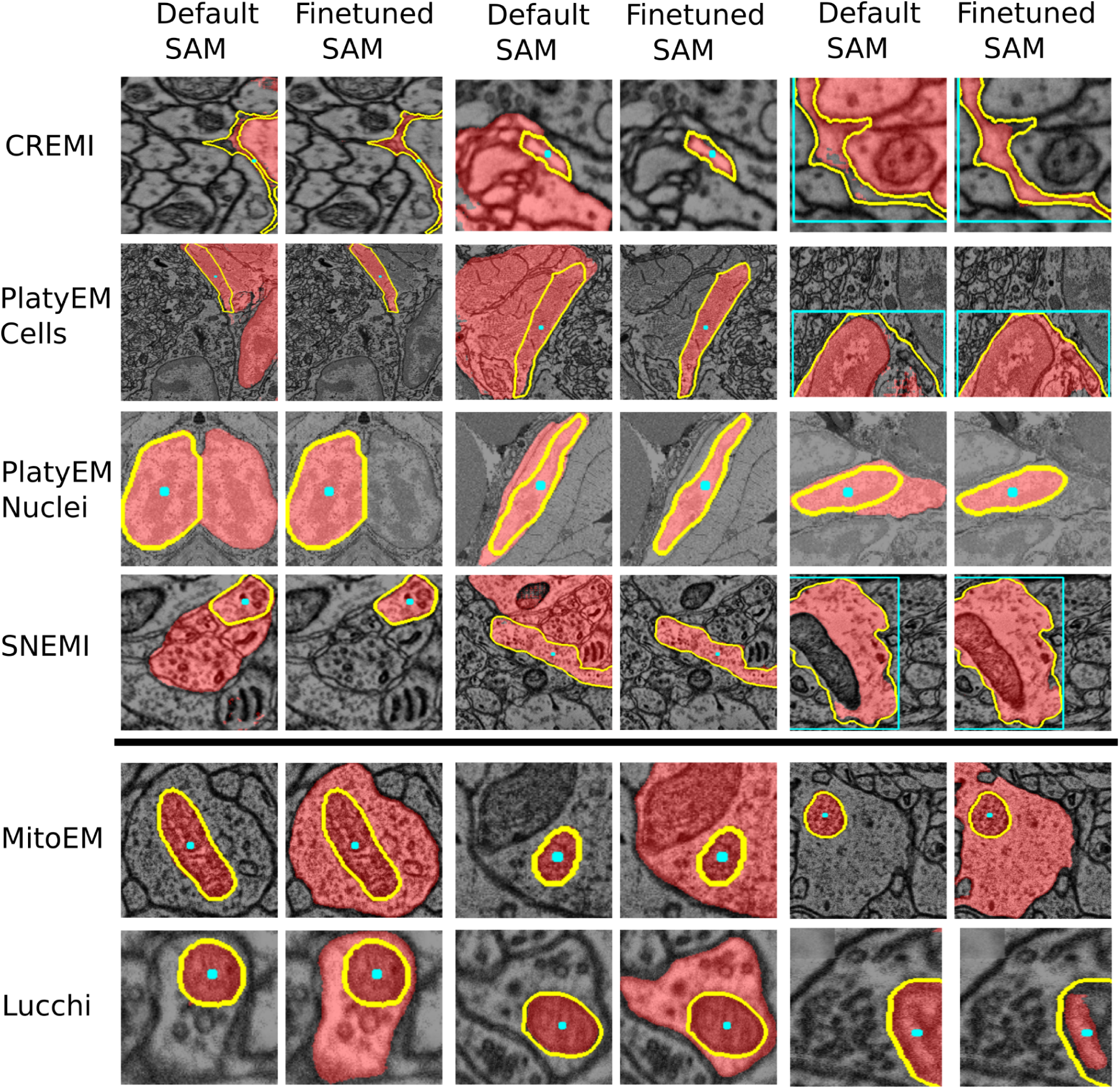
Qualitative comparison of finetuned EM and default model. The images above the bold line were generated by selecting objects with the highest improvement of IOU score of the finetuned compared to the default model, the images below using the opposite approach. Here, we see that the generalist model improves segmentation performance in the case of point and box annotations for segmenting neurons or cells by segmenting only single objects instead of groups, and yielding better segmentations for objects of complex shape or objects with internal structure, e.g. due to organelles. Conversely the segmentation performance is worse for mitochondria because the model segments the full surrounding cell / cellular compartment instead of the mitochondrion. These observation support our finding that training a single EM generalist model is not feasible, but that improved models for specific structures can be trained.

**Supplementary Figure 5:**
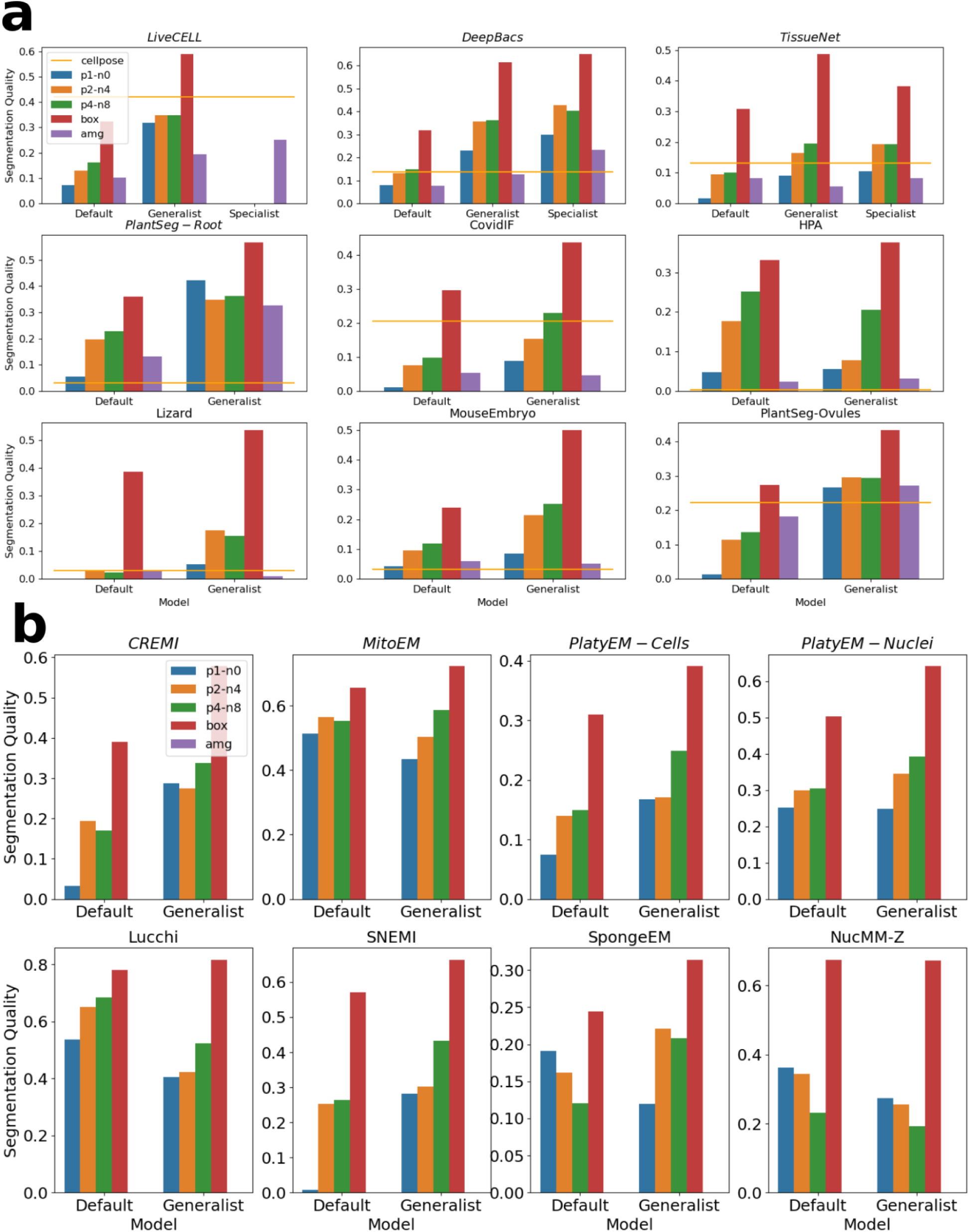
Comparison of the generalist / finetuned ViT-B models for LM (**a**) and EM (**b**). Overall these results show the same trends as for the ViT-H models (Figure 3 and 4), with generally lower performance for the ViT-B models.

**Supplementary Figure 6:**
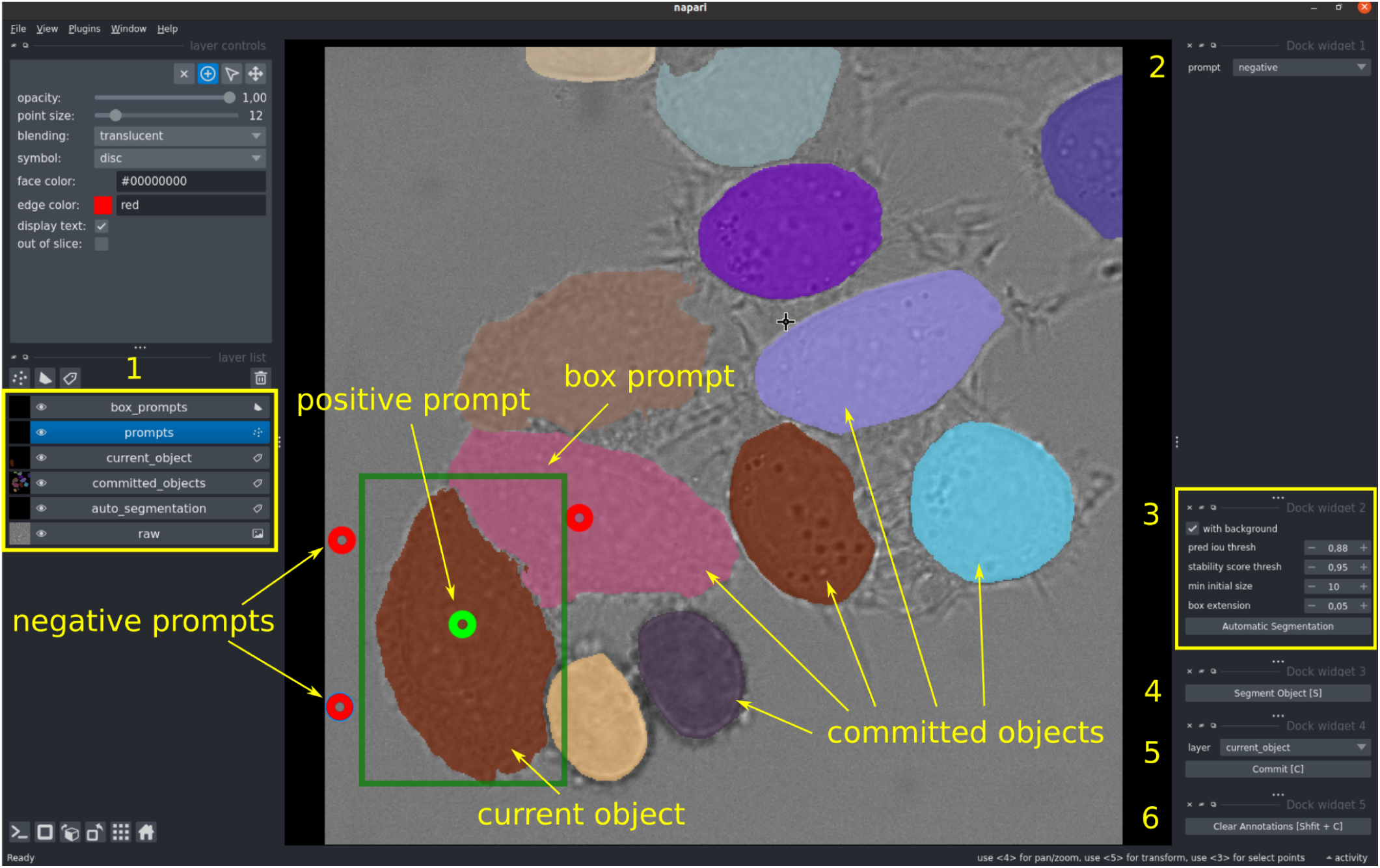
User interface of the 2d annotation tool.

### Footnotes

1 Some LM label-free imaging techniques can also unspecifically label cellular components. The datasets studied in LM however contain either imaging with specific resolution or are label-free but of low resolution so that subcellular structures cannot be resolved.

2 https://github.com/constantinpape/torch-em

